# Edaphic controls on genome size and GC content of bacteria in soil microbial communities

**DOI:** 10.1101/2021.11.17.469016

**Authors:** Peter F. Chuckran, Cody Flagg, Jeffrey Propster, William A. Rutherford, Ella Sieradzki, Steven J. Blazewicz, Bruce Hungate, Jennifer Pett-Ridge, Egbert Schwartz, Paul Dijkstra

## Abstract

Bacteria in soil microbial communities are crucial to terrestrial ecosystem function, yet our understanding of the fundamental characteristics of their genomes, such as GC content and genome size, is not complete. Much of our understanding of the mechanisms which shape their genomic traits is derived from other systems or from isolated bacteria. Here we determined average genome size, GC content, codon usage, and amino acid content from 398 soil metagenomes across a broad geographic range and used machine-learning to determine which environmental parameters most strongly explain the distribution of these traits. We found that genomic trait averages were most related to pH, which we suggest is primarily due to the correlation of pH with several environmental parameters, especially soil carbon content. Low pH soils had higher carbon to nitrogen ratios (C:N) and tended to have communities with lower GC content and larger genomes, potentially a response to increased physiological stress and a requirement for metabolic diversity. Conversely, smaller genomes with high GC content were associated with high pH and low soil C:N, indicating potential resource driven selection against AT base pairs. As soil bacteria tend to be more carbon limited, smaller genomes with higher GC content may reduce the cost of reproduction in carbon-limited soils. Similarly, we found that nutrient conservation also applied to amino acid stoichiometry, where bacteria in soils with low C:N ratios tended to code for amino acids with lower C:N. Together, these relationships point towards fundamental mechanisms that underpin nucleotide and amino acid selection in soil bacterial communities.

## MAIN TEXT

In bacteria, nutrient constraints exert influence on traits such as genome size, GC content, codon frequency, and amino acid content [1–3]. For free-living bacteria, low nutrient concentrations often select for genomic traits that reduce the cost of reproduction, such as low GC content and smaller genomes [3]. A GC base pair requires more energy to produce than an AT base pair [4, 5]. Moreover, since the GC base pair has a carbon to nitrogen ratio (C:N) of 9:8 (1.13), whereas a AT base pair has a C:N of 10:7 (1.42), GC-rich genomes require more nitrogen, which may be disadvantageous in nitrogen-limited environments. Additionally, there is a correlation between the elemental requirements for nucleic and amino acids, such that codons with higher GC content (which have lower C:N) tend to code for amino acids with low C:N [6]. Therefore, nucleotide frequency not only controls the energy requirements for DNA synthesis, but also directly relates to the nutritional demand for amino acid synthesis [7].

However, much of the existing and foundational literature on processes controlling genomic traits in free-living bacteria are based on the study of marine isolates [8] and aspects of this framework may not cleanly transpose onto soil bacteria. For example, we had previously found that community-averaged GC content and genome size were positively correlated among marine metagenomes—a relationship attributed to N-limitation; however, the correlation between GC content and genome size was negatively correlated between metagenomes collected from soils [9]. Since the growth of soil bacteria is thought to be more limited by carbon than nitrogen [10, 11], we hypothesized that the distribution of genomic traits in soil bacteria might exhibit unique patterns reflecting carbon limitation. Specifically, because the GC base pair (1.13) has a lower C:N than the AT base pair (1.42), we predicted communities would exhibit higher GC content and smaller genomes when carbon availability is low. Carbon limitation has been indicated as a potential driver of high GC content in microbial models [12], and previous studies have shown that microbial communities from bulk soil (often carbon-poor) tend to have smaller genomes and higher GC content than those in the rhizosphere [13]. However, the relationship between soil carbon and the genomic traits of soil bacteria has not been properly assessed and represents a fundamental unknown in our understanding of belowground microbial life. To better understand the relationship between genomic traits of soil bacteria and environmental characteristics, we analyzed 398 metagenomes collected and sequenced by the US-based National Ecological Observation Network (NEON) [14] across a broad geographic scale and analyzed community-averaged genomic traits alongside a range of environmental and edaphic properties.

## RESULTS AND DISCUSSION

We hypothesized that microbial communities in low carbon environments would exhibit smaller genome sizes, higher GC content, and amino acid composition with a lower carbon to nitrogen ratio (C:N); which would be predominantly driven by carbon limitation in the bacterial community. To test this, we assessed the relationship between extractable soil C:N (C_extr_:N_extr_) and genomic traits across all NEON sites (Fig. 1A). We found a negative correlation between GC content and C_extr_:N_extr_ (*p* < 0.001; Fig. 1B), and a positive correlation between C_extr_:N_extr_ and both estimates of average genome size (*p* < 0.001; Fig. 1C). Further, we found that average genome size and GC content calculated from bacterial contigs were negatively correlated (*p* < 0.01, Fig. 1D). This relationship is unique in comparison to free-living marine bacteria, where smaller genomes tend to have lower GC content [3].

**Figure 1:**
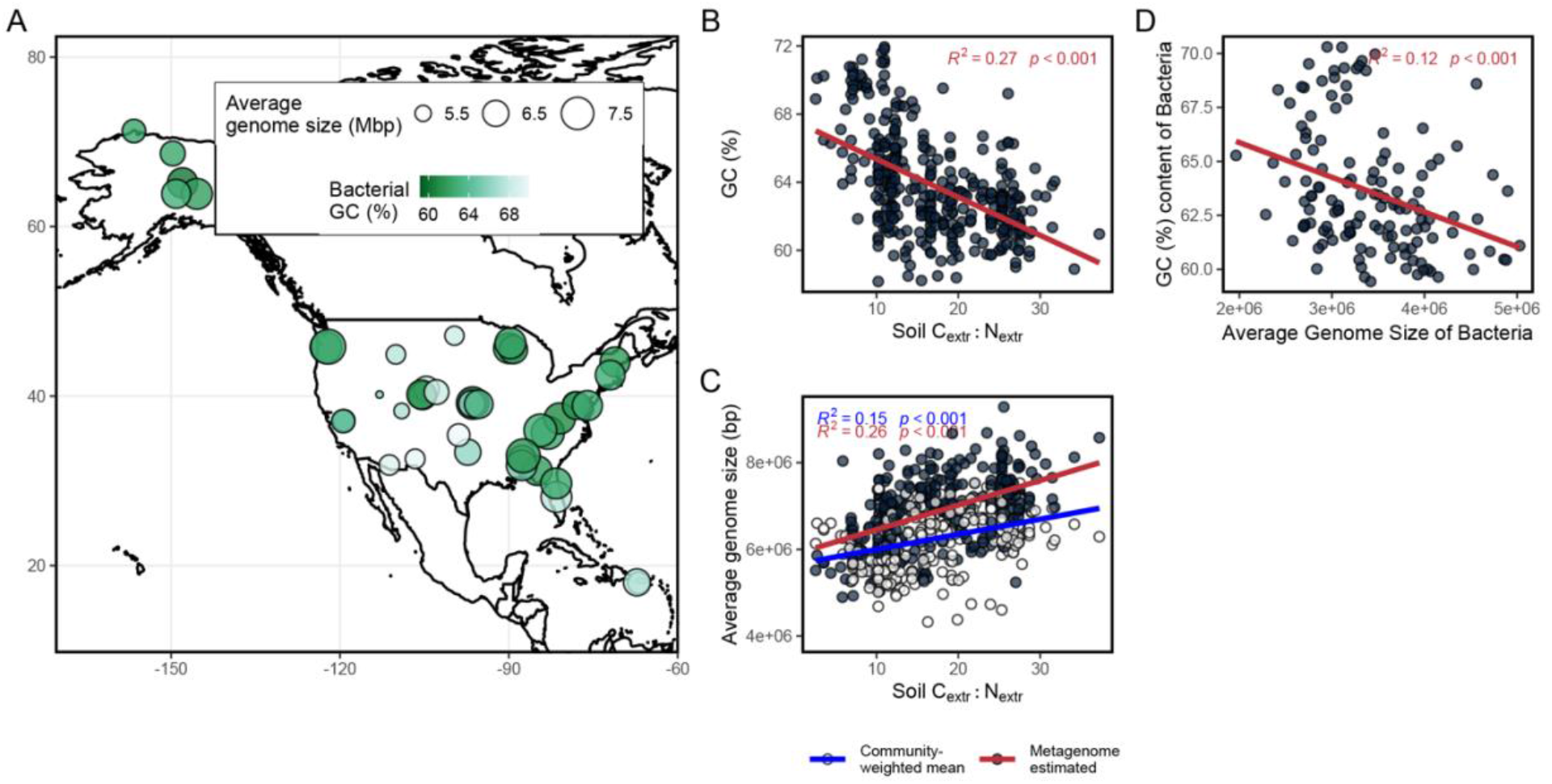
Distribution of genomic traits across sites and soil extractable carbon and extractable nitrogen ratios (C_extr_:N_extr_); **(A)** Geographic distribution of sites, with mean bacterial GC and estimated average genome size. **(B)** Relationship between C_extr_:N_extr_ and bacterial GC content (%) (linear regression, *p* < 0.001). **(C)** Relationship between C_extr_:N_extr_ and genome size, estimated from the number of single copy genes per metagenome (dark circles, linear regression, *p* < 0.001) and from 16S rRNA gene datasets and genome size estimated from isolates (open circles, linear regression, *p* < 0.001). **(D)** The average GC content of all bacterial contigs in a metagenome vs average genome size estimated from bacterial contigs.

The C:N of the sum of all predicted amino acids was positively correlated with soil C_extr_:N_extr_ (*R*^2^=0.24, *p* < 0.001, Fig. 2A), and closely tracked metagenome GC content (*R*^2^=0.51, *p* < 0.001, Fig. 2B), reflecting the resource alignment between the stoichiometry of nucleic acids in codons and their corresponding amino acids [6]. We found that synonymous codon usage skewed towards codons with a higher GC content, perhaps best represented by the strong preference for guanine and cytosine at fourfold degenerate sites (*p* < 0.01; Fig. 2C). This suggests that GC content is not solely a consequence of amino acid composition, and that there is additional nucleotide bias. Notably, we also observed a higher abundance of cytosine than guanine and fourfold degenerate sites (Fig. 2C) that could be a consequence of the lower metabolic cost of cytosine production [4]. Preferential selection for codons with higher AT content was most pronounced where soil C:N was high. For each amino acid, codons with higher GC were more often negatively correlated with soil C_extr_: N_extr_ compared to codons with lower GC, which more often were positively correlated with soil C_extr_:N_extr_ (Fig. 2D&E).

**Figure 2:**
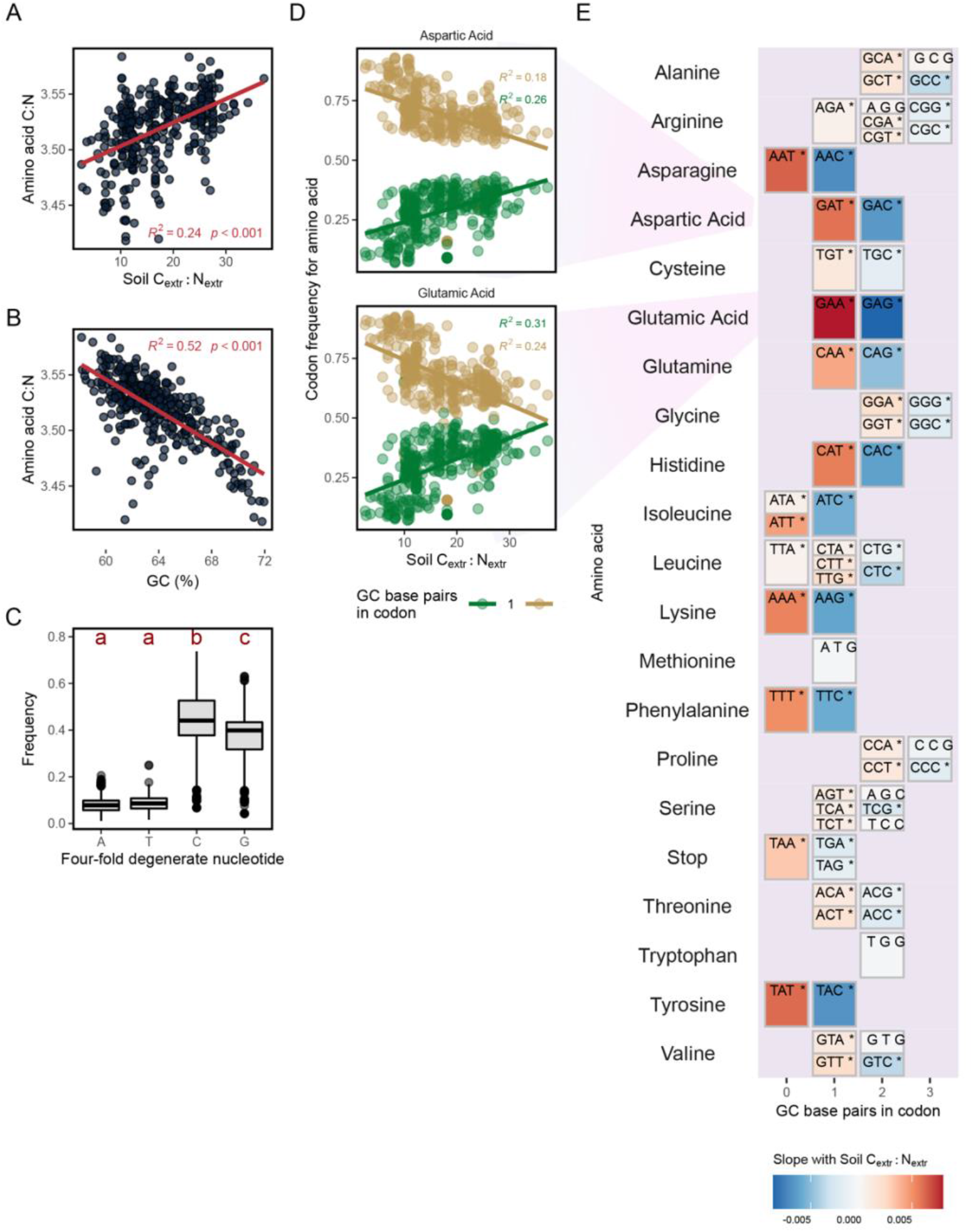
Averaged nucleotide, codon, and amino acid composition of bacteria in metagenomes **(A)** Relationship between C_extr_:N_extr_ and bacterial amino acid C:N ratios averaged per metagenome (linear regression, *p* < 0.001). **(B)** Relationship between bacterial GC and pooled amino acid C:N ratios (linear regression, *p* < 0.001). **(C)** The distribution of nucleotides at the third position in fourfold degenerate codons across all metagenomes, with letters corresponding to groups identified via Tukey’s post-hoc test (*p* < 0.01). **(D)** The relationship between codon frequency and soil C_extr_:N_extr_ shown for aspartic acid and glutamic acid, with number of GC base pairs in each codon being indicated by color (linear regression, *p* < 0.001). **(E)** The relationship between codon frequency and soil C_extr_:N_extr_ shown for each codon, with color indicating the slope of the relationship and an asterisk indicating significance (linear regression, *p* < 0.05). Codons are arranged left to right in increasing number of GC base pairs.

In line with our original hypothesis, lower soil C:N was associated with communities averaging smaller genomes, higher GC, and lower C:N of amino acids. However, genomic traits in soil microbial communities may also be driven by other environmental factors, such as temperature [15, 16] and pH [17]. To assess the relationships between genomic traits and other environmental drivers, we used a machine learning, random-forest model approach to determine the environmental variables that explain the most variance in GC content and predicted average genome size. With this model, we assessed the importance of over 100 environmental factors and geographic range in shaping genomic features.

Random forest models indicated that GC content and the average genome size of a community were most strongly related to soil pH (Fig. 3A), where soils with low pH fostered communities with low GC content (Fig. 3B) and larger average genome size (Fig. 3C). Large fungal genomes tended to be more present in low pH soils (Supplemental Fig. 1A) and correlated with larger community-average genome size (Supplemental Fig. 1B). However, similar relationships were observed when genome size was predicted from single copy genes detected in bacterial contigs (Supplemental Results & Discussion; Supplemental Fig. 1C) as well as 16S rRNA gene taxonomy (Supplemental Fig. 2), indicating that the observed trends are also being driven by changes in bacterial genome size (as opposed to solely changes in the abundance of non-bacterial reads).

**Figure 3:**
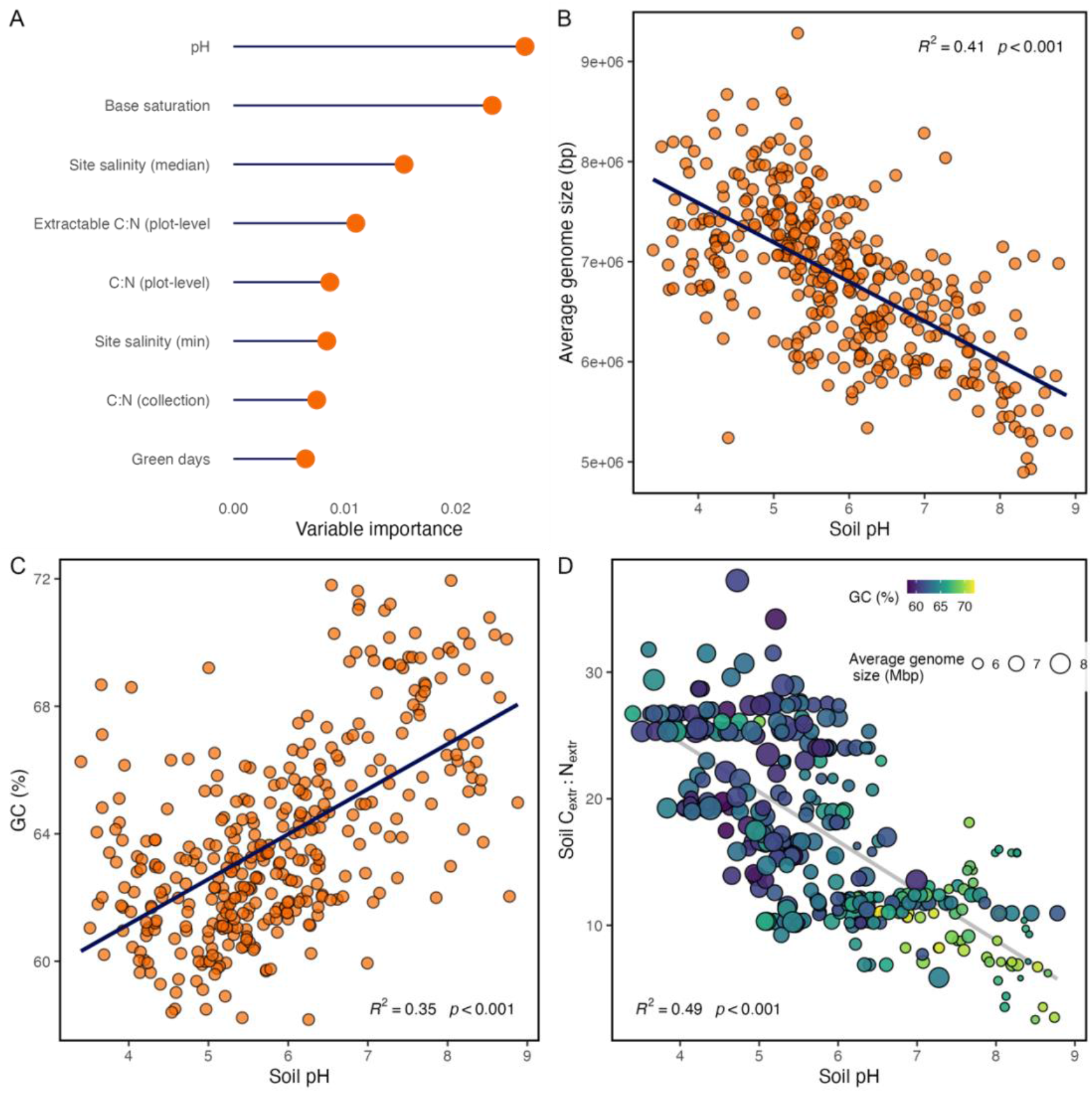
Results from the random forest model and relationships between soil pH and genomic traits; **(A)** Variable importance plot for the top 8 environmental parameters predicting GC content from the random forest model (RMSE = 0.017; *R^2^* = 0.66). **(B)** The relationship between soil pH and GC content (%) of bacterial contigs, with a linear relationship as selected by the random forest model (*p* < 0.01).**(C)** The relationship between soil pH and average genome size (derived from metagenomes) with a linear relationship as selected by the random forest model (blue; *p* < 0.01) **(D)** The relationship between soil pH and soil C_extr_:N_extr_ with points colored by bacterial GC content and point size corresponding to average genome size.

There are several reasons why pH might be correlated with this set of genomic characteristics in soil bacteria. Soil pH represents the intersection of numerous environmental vectors and, accordingly, we hypothesize that there are several mechanisms underpinning the relationship between pH and genomic traits. First, low pH causes physiological stress in soil bacteria and is often associated with a greater number of repair mechanisms, such as chaperones [18]. This might preferentially select for bacteria with a greater investment in stress alleviation and maintenance, and thus larger genomes. Second, low pH is often associated with the accumulation of soil organic carbon (SOC). Since soil pH is largely driven by the balance between precipitation and evapotranspiration [19], low pH often coincides with greater precipitation excess and primary production. Higher biomass inputs into acidic soils, combined with a reduction in the decomposition rate due to low pH, results in the build-up of SOC [18]. The accumulation of SOC not only alleviates carbon limitation—which may reduce GC content—but also potentially favors larger genomes with increased metabolic diversity. It has been suggested that the requirement for increased metabolic diversity might explain why soil bacterial genomes tend to have large genomes [20] and, similarly, we found that soils with lower pH were associated with higher C_extr_:N_extr_ as well as larger genomes and lower GC content (Fig. 3D). High pH in soil can also result in the build-up of microbial biomass and increases in SOC [18], which could explain why we found that more alkaline soils had higher C_extr_:N_extr_, larger genomes, and lower GC content than soils with a neutral pH. Third, genomic traits in soil bacteria may relate to other forms of stress coinciding with pH. Aridity has been shown to drive streamlining in certain soil bacteria [21] and, as discussed above, influences the pH in soil. Previous work has shown relationships between precipitation, pH, and genome size [17], and in a previous analysis we found that soil metagenomes collected from both hot and cold deserts often had smaller genomes and greater GC content than soils collected in more mesic systems [9]. Although we did find a positive relationship between mean annual precipitation and average genomes size (Supplemental Fig. 3), the relationship was not as strong as that for pH or soil C_extr_:N_extr_.

Our work demonstrates that soil pH determines the broad-scale distribution of genomic traits between soil bacterial communities, which we suggest can be attributed to pH being a metric that captures multiple environmental parameters, such as soil nutrients and precipitation patterns, as well as physiological stress. Soil pH is well established driver of community composition in soils [22] and our results emphasize the degree to which pH dictates belowground microbial life and perhaps influences the evolution of soil bacteria. Additionally, we found several trends that suggest that selection pressure in soil bacterial communities might reflect carbon limitation; for example, the negative relationship between genome size and GC content. The overall influence of C:N on nucleotide selection is similar to what has been observed in marine systems (i.e. low C:N selects for higher GC content, and higher C:N selects for lower GC content); however, the reduction in genome size with high GC content and low soil carbon is distinct, and suggests that carbon limitation is driving the distribution of these traits. These results are derived from community averages and more work must be done to uncover both the mechanisms and taxonomic level where the observed changes in genomic traits occur. However, it is evident that the distribution of genomic traits in soil is related to edaphic properties and deserves further study.

## METHODS

Data for this project was gathered from the National Ecological Observation Network (NEON)— an observational network collecting ecological data from across the United States, funded by the US National Science Foundation. NEON maintains 81 field sites in a number of distinct biomes across the US. Terrestrial sites include a central meteorological station as well as many dispersed plots from which samples are collected [23]. A full description of NEON sites can be found at https://www.neonscience.org/field-sites/about-field-sites. Data products used in this analysis can be found in Supplemental Table 1. A full list of data used in this analysis, as well as associated descriptions of this data, can be found in Supplemental Tables 2 and 3, and metagenome assembly statistics in Supplemental Table 4.

### Metagenomic traits

From the NEON data portal (https://www.neonscience.org/data), we accessed the metadata for all available metagenomes as of January 2021. We selected 398 metagenomes which demonstrated the greatest read depth while maximizing the number of collection sites (43 total). In January of 2021, selected metagenomes were downloaded from the NEON data portal to the high-performance computing cluster at Northern Arizona University.

Raw reads were QC filtered using Trimmomatic v0.39 (parameters: TruSeq2-PE.fa:2:30:10 LEADING:10 TRAILING:10 SLIDINGWINDOW:10:20 MINLEN:50) [24]. We then used the program MicrobeCensus [25] to determine the average genome size of a microbial community from the QC-filtered reads. MicrobeCensus estimates the total number of single copy genes to create an estimate of average genome size from unassembled reads. Contigs were assembled from QC-filtered reads using MEGAHIT v1.2.9 (parameters: --k-list 21,29,39,59,79 --min-contig-len 400) [26] and read depth for each contig was determined using BBMap v38.87 (bbmap.sh default parameters) [27]. Open reading frames (ORFs) were then identified using Prodigal v2.6.3 (parameters: -p meta) [28] and taxonomy was assigned to each ORF using Kaiju v1.7.4 using the default parameters against a NCBI *nr* database (constructed with: kaiju-makedb -s nr_euk) [29]. HMMER was used to annotate genes against the pfam database [30].

Using the read depth for each contig and the estimated taxonomic identity for each ORF, we calculated a depth-adjusted GC content for the bacterial contigs in a metagenome. Similarly, we calculated the depth-adjusted amino acid content for each metagenome. Using known chemical formulas for each amino acid and the depth-adjusted amino acid content, we calculated an estimate of the total C:N ratio of the amino acid pool in each metagenome. For amino acids with fourfold degenerative sites (alanine, glycine, proline, threonine, valine), we calculated the frequency of nucleotides at the third position and averaged these values across the selected amino acids. This provided a metagenome-level estimate of nucleotide frequencies for fourfold degenerative sites.

Since our estimates of average genome size for a metagenome was based on the QC filtered reads, non-bacterial reads could potentially bias these estimates. To assess whether the trends we observed were driven by changes in bacterial average genome size, we generated another estimate of genome size based solely on the assembled bacterial contigs. We searched our bacterial pfam gene annotations for 139 single copy genes (based on Campbell et al. 2013 [31]) and adjusted for coverage and read depth. We then used the mean number of single copy genes as an estimate of the number of bacterial genomes represented in our metagenomic assembly. We divided the total number of base pairs of our bacterial contigs (adjusted for read depth) by the estimated number of bacterial genomes to generate an estimate of bacterial average genome size. Metagenomes with fewer than 200 million assembled bacterial base pairs where discarded—which provided a similar sample size to MicrobeCensus with default parameters.

Specific commands for metagenomic assembly, mapping, and annotation, as well as custom scripts for genetic data processing, can be found at the following Github repository page: https://github.com/PChuckran/Soil_genomic_traits/tree/main/code/upstream_bioinformatics.

### Traits from 16S rRNA genes

We also estimated average genome size using a community-weighted mean derived from aligning bacterial 16S rRNA gene sequences against a database of genomes of known size. 16S rRNA gene amplicon datasets were downloaded from the NEON data portal. We selected samples from the same sites and collection times as the metagenomes used in this study. To build a reference database, in August 2021 we downloaded all available bacterial 16S gene sequences and associated metadata from the Genome Taxonomy Database (release 202) [32] and created a BLAST [33] database (parameters: makeblastdb -parse_seqids -dbtype nucl) consisting of high quality (>95% complete, <5% redundant) bacterial genomes. Operational taxonomic units (OTUs; clustered at 97%) identified by NEON were then aligned to our high-quality database using BLAST [33] (parameters: blastn -max_target_seqs 1). The best alignment (at >99% identity) was used as the assigned taxonomy. The genome size of each assigned OTU was then adjusted for read depth and used to calculate a community-weighted mean genome size with mean coverage (read depth) as the weight factor. This provided us with a secondary estimate of genome size based only on the bacterial community. Although genome size for a given taxon can vary, we believe this approach still detects general trends in genome size with changes in community composition. Importantly, the source of error in this estimate is different from the metagenome-derived estimate (i.e. unknown intraspecific variations as opposed to the presence of non-bacterial genomes). We also used the GC content of the 16S rRNA gene itself to generate an additional estimate of GC content. Similar trends of genomic traits with edaphic characteristics would provide independent support that these relationships are not due to errors in the methods used for estimation.

### Analysis with environmental characteristics

From the NEON data portal and meteorological data, we gathered 205 environmental parameters associated with each site and date of collection. Data were accessed between January and April 2021. Where possible, edaphic characteristics were paired with metagenomes from the same soil sample. Otherwise, metagenomes were paired with data from plot or site level averages. Microbial PLFA biomarkers were used to calculate fungal to bacterial biomass ratios [34, 35]. Standard precipitation and standard precipitation evapotranspiration indices (SPI and SPEI) were gathered from GRIDMET [36] using Google Earth Engine [37]. Linear regression analysis was used to determine the relationship between soil carbon and genomic traits. All models were constructed in R (v 3.6.1) [38].

Random forest models were constructed using the Tidymodels [39] and ranger [40] packages in R, with the objective of determining the environmental parameters which most strongly influenced genomic traits. A non-parametric, machine learning/random forest regression model approach was selected given its proven ability to handle non-linear, complex interactions in space and time across multiple predictor variables [41]. Prior to tuning the random forest hyperparameters, predictors which did not provide data for at least two-thirds of metagenomes were dropped, and the remaining variables were scaled and centered prior to analysis. The full dataset contained several environmental measurements which were either very similar or functionally identical metrics collected at multiple levels (e.g., site, plot, and soil core). Redundant predictors were removed by first running a random forest which included all available predictors, which were then ranked by variable importance. We calculated Pearson’s correlation coefficients between all predictors and then removed those which were highly correlated (R^2^ > 0.8) with a predictor which explained more variability. We then ran a random forest model which included only remaining predictors.

The strongest predictor of genomic traits, as indicated by random forest models, was then fit to the full dataset. It is important to note that the random sampling associated with this approach means that the best-fit relationship indicated by the random forest, although potentially more accurate as to the true nature of the relationship, may not represent the best-fit for the entire dataset. The code for custom bioinformatics analysis, downstream statistical analysis, and refined data is publicly available at https://github.com/PChuckran/Soil_genomic_traits.

## ACKNOWLEDGEMENTS

We would like to thank Anita Antoninka, Michaela Hayer, Alicia Purcell, Junhui Li, Megan Foley, Raina Fitzpatrick, Bram Stone, Victoria Monsaint-Queeney, and Carl Roybal for their intellectual contributions to this work.

