## Supplemental Material for "Edaphic controls on genome size and GC content of bacteria in soil microbial communities"

SUPPLEMENTAL RESULTS AND DISCUSSION

***Alternate estimates of genomic traits***

Although the average genome size of a community as predicted by single copy genes could be influenced by the abundance of nonbacterial reads—such as fungi, which are highly abundant in low pH soils—we found a similar relationship when we predicted genome size using single copy genes detected in the assembled bacterial contigs (n = 133, *R*^2^=0.14, *p* < 0.01, Supplemental Fig. 1). These estimates for genome size were smaller than those from the QC-filtered reads, which we attribute to the fact that well-known single copy genes are more likely to be annotated. Still, the relationship between pH and this estimate of genome size indicates that the average genome size of the bacterial population is increasing with low pH.

We also found a relationship between pH and average genome size for bacterial species when genome size was predicted by aligning 16S rRNA gene sequences with a database of bacterial isolates of known size (Supplemental Fig. 2A). Estimates of average genome size predicted from metagenomic and 16S rRNA genes were also correlated (*R^2^*=0.3*, p* < 0.01), although estimates were substantially smaller than metagenome-based estimates (Supplemental Fig. 2B). We found that the difference between these estimates was not related to fungal biomass (linear regression of %difference in genome size estimates ~ fungal:bacterial biomass (PLFA); *R^2^* = 0.01, *p* > 0.05), suggesting that this bias is unlikely driven solely by large fungal genomes in the metagenomes. One potential explanation is that bacterial genome size of a given taxa in soil may be larger than what is recorded in genomic databases. Soil genomes tend to be larger than in other systems [1–3], and since representatives from GTDB can represent a consensus or composite from multiple environments, identity-based estimates could potentially underestimate genome size. Ultimately, it is not possible to get at the source of this error from the data presented here. However, the correlation between these estimates, and their general agreement with respect to changing edaphic properties, increases our confidence that the observed trends with environmental variables reflect real-world phenomena. Similarly, we found that the GC content of the 16S rRNA genes themselves followed a similar patter to those of the bacterial reads of the metagenomes with respect to edaphic properties as the GC content from the metagenomes (Supplemental Fig. 2C & 2D). As with the estimate for genome size, the coincidence of these trends supports our assessment that environmental properties are shaping the genomic traits of soil bacteria.

SUPPLEMENTAL TABLES AND FIGURES

**Supplemental Table 1:**

NEON data products used in this study and their associated materials.

| **Data product ID** | **Data product name** | **doi** | **Date accessed** |
| --- | --- | --- | --- |
| DP1.10086.001 | Soil physical and chemical properties, periodic | https://doi.org/10.48443/3qtw-w090 | 25-Feb-21 |
| DP1.10107.001 | Soil microbe metagenome sequences | https://doi.org/10.48443/fzzj-g053 | 25-Feb-21 |
| DP1.10109.001 | Soil microbe group abundances | https://doi.org/10.48443/3qkb-3m53 | 25-Feb-21 |
| DP1.10104.001 | Soil microbe biomass | https://doi.org/10.48443/xnbn-rw33 | 25-Feb-21 |
| DP1.10108.001 | Soil microbe marker gene sequences | https://doi.org/10.48443/ybrs-zv89 | 25-Feb-21 |
| DP1.10023.001 | Herbaceous clip harvest | https://doi.org/10.48443/xjxw-2p18 | 25-Feb-21 |
| DP1.10033.001 | Litterfall and fine woody debris production and chemistry | https://doi.org/10.48443/jvrg-xp36 | 25-Feb-21 |
| DP1.10067.001 | Root biomass and chemistry, periodic | https://doi.org/10.48443/5wvb-ww20 | 25-Feb-21 |
| DP1.10111.001 | Site management and event reporting | https://doi.org/10.48443/dtjh-q277 | 25-Feb-21 |
| DP1.10047.001 | Soil physical and chemical properties, distributed initial characterization | https://doi.org/10.48443/x9aj-3647 | 25-Feb-21 |
| DP1.00096.001 | Soil physical and chemical properties, Megapit | https://doi.org/10.48443/rfmw-p030 | 25-Feb-21 |

**Supplemental Table 2:**

All the summarized data used in this analysis. Please refer to separate csv file titled Supplemental_table_2.csv

**Supplemental Table 3:**

A description of the columns in Supplemental Table 2. Includes the level at which the data was collected (plot, site, sample), description, and relevant notes. Please refer to attached xlsx file Supplemental_table_3.xlsx

**Supplemental Table 4**

Summary statistics for metagenome assemblies. Regrettably, some of the assembly statistics were lost due to a file management system error; however, the provided data indicate the range of assembly quality for this study. Please see Supplemental_table_4.csv

**Supplemental Figure 1**

The influence of fungi on estimations of genome size with pH. **(A)** The fungal to bacterial ratio as determined by PLFA analysis as it relates to soil pH; **(B)** The relationship between average genome size and fungal to bacterial ratios (PLFA); and **(C)** the relationship between pH and average genome size determined from the assembled bacterial contigs.

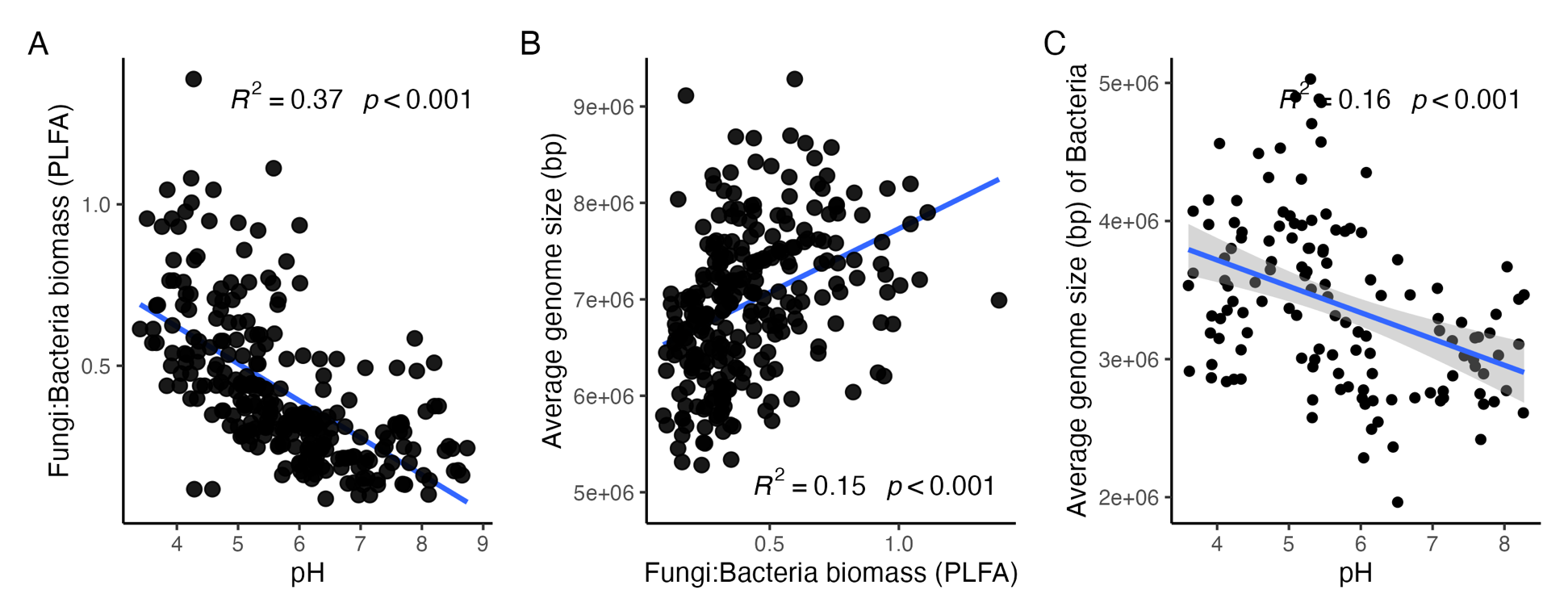

**Supplemental Figure 2:**

A comparison of taxonomy based estimates of genomic traits predicted from 16S rRNA genes. (**A**) Average genome size predicted from taxonomy as a function of pH. (**B**) The relationship between metagenome-based estimates of average genome size and average genome size predicted from taxonomy. The relationship between the mean GC content (%) of 16S rRNA genes and soil pH (**C**) and extractable soil C:N (**D**)

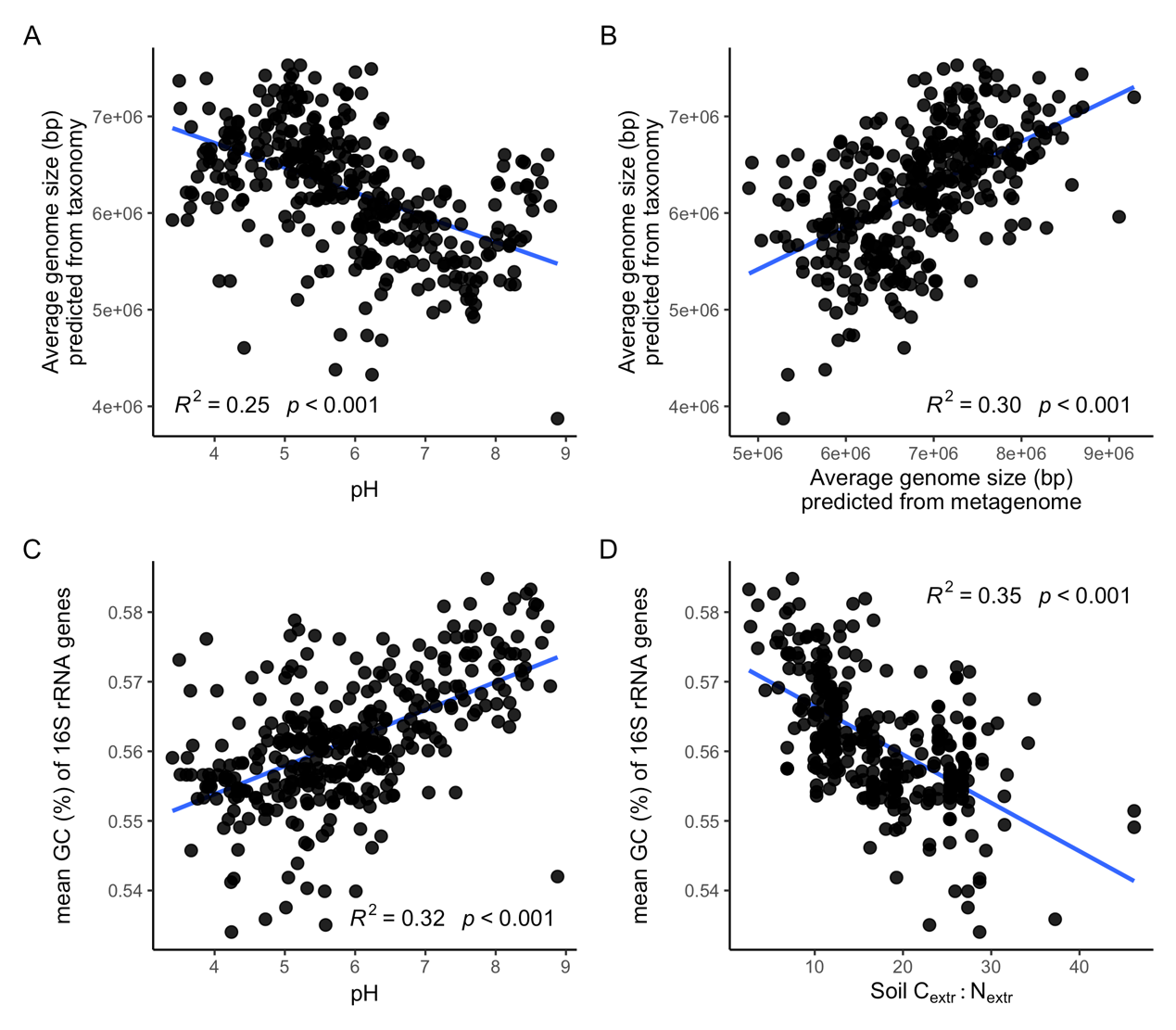

**Supplemental Figure 3:**

The relationship between mean site-level measurements of mean annual precipitation (mm) and average genome size.

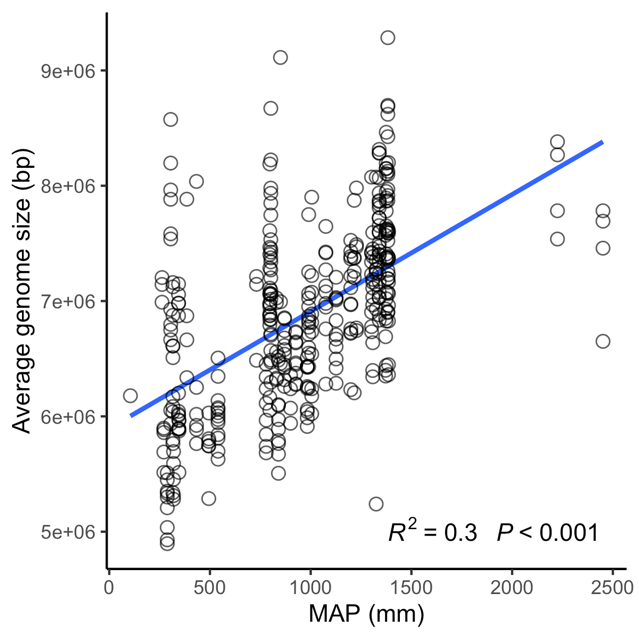
